# METTL14-dependent m^6^A modification restrains interferon signaling to prevent myocarditis and dilated Cardiomyopathy

**DOI:** 10.64898/2026.04.02.716218

**Authors:** Yu Xi, Jacob Kuempel, Sukwon Choi, Patrick DeSpain, Tianru Zhang, Jiaying Zhu, Abi Osborn, Reed Rivera, Songxiao Zhong, Xiaoyang Wu, Yong-Xiao Wang, Zhenyu Li, A. Philip West, Changhao Li, Carl W. Tong, Xiuren Zhang, Xu Peng

## Abstract

The impact of inflammation on heart failure is increasingly recognized; but how cardiomyocyte restrains innate immune activation remains poorly defined, and nor does the role of N⁶-methyladenosine (m⁶A) modification in maintaining cardiac immune homeostasis. Here, we demonstrate that cardiomyocyte-specific deletion of the m⁶A methyltransferase METTL14 triggers myocarditis, dilated cardiomyopathy, and premature lethality. Meanwhile, widespread hypomethylation and upregulation of innate immune and necroptosis-related transcripts in *Mettl14*-deficient hearts exemplified by IFN-1 and STAT1. Mechanistically, METTL14 deficiency promotes RIPK1 accumulation thereby priming cardiomyocytes for necroptosis and inflammatory cell death. Genetic ablation of IFN-I receptor *Ifnar1* can largely rescue the processes and improve cardiac function and survival. Furthermore, METTL14 loss disrupts mitochondrial integrity and autophagy/mitophagy flux, suggesting mitochondrial dysfunction–driven innate immune activation upstream of IFN-I signaling. Collectively, these findings identify METTL14-mediated m⁶A modification as a critical safeguard against cardiomyocyte-intrinsic IFN-I signaling and necroptosis and establish an epitranscriptomic–innate immune axis that drives inflammatory heart failure.

## Introduction

Heart failure is a rapidly growing global health challenge, rising in prevalence from 5 million cases in 2012 to 6.7 million in 2023 in the United States and affecting more than 64 million individuals worldwide in 2019 (1, 2). The COVID-19 pandemic further exacerbates this burden, with approximately 11.6 new heart failure diagnoses per 1,000 SARS-CoV-2 infections (3). Inflammation is an independent predictor of adverse outcomes in heart failure. A large body of clinical and experimental evidence has now implicated inflammation as a key pathogenic driver (4, 5). Despite the strong association between systemic inflammation and heart failure progression, clinical trials targeting classic inflammatory mediators, exemplified by tumor necrosis factor-α and interleukins, have produced disappointing results (5). These findings highlight the urgent need to uncover heart-specific inflammatory mechanisms for heart failure and thus may provide new therapeutic opportunities.

Autoimmune myocarditis is a major cause of non-ischemic heart failure and is characterized by immune-mediated cardiomyocyte injury that frequently progresses to dilated cardiomyopathy (6). The introduction of immune checkpoint inhibitors (ICIs) has revolutionized cancer therapy but also revealed fulminant myocarditis as a severe immune-related adverse event (7). In ICI-induced myocarditis, excessive T-cell activation and interferon-driven inflammation converge to breach cardiac immune tolerance, mirroring key pathological features of idiopathic autoimmune myocarditis (7, 8). Yet, the identity of cardiomyocyte-intrinsic signaling pathways that contribute to interferon-mediated myocarditis and its progression toward dilated cardiomyopathy and underlying mechanism are largely unknown.

Type I interferons (IFN-I), primarily IFN-α and IFN-β, are essential antiviral cytokines but are increasingly recognized as potent drivers of non-infectious cardiac injury (9). Chronic or excessive IFN-I activation promotes inflammation, adverse remodeling, and systolic dysfunction, ultimately contributing to heart failure (10, 11). Dysregulated expression of IFN-I–related genes is strongly associated with distinct heart failure subtypes, suggesting that interferon signaling contributes to the heterogeneity of cardiac dysfunction (12). Following myocardial infarction (MI), the cells in the border zone can activate an interferon regulatory factor 3 (IRF3)-dependent IFN-I program that impairs fibroblast-mediated repair and exacerbates ventricular dysfunction and rupture (10). Genetic models with defective nucleic acid clearance, such as Trex1 knockout mice, develop lethal autoimmune myocarditis and dilated cardiomyopathy via cyclic GMP-AMP synthase-stimulator of interferon genes (cGAS–STING)–IFN-I signaling (13, 14). Most recently, the Z-DNA/RNA sensor ZBP1 has been shown to activate IFN-I signaling and drive inflammatory cardiomyopathy during mitochondrial stress, linking nucleic acid sensing, necroptosis, and heart failure (11). Furthermore, nuclear envelope disruption causes genomic DNA to leak into the cytosol, thus activating the cGAS–STING–IFN-I pathway and accelerating dilated cardiomyopathy progression (15). Collectively, these studies position chronic or sterile IFN-I activation as a central pathogenic axis that bridges nucleic-acid-triggered innate immune activation to progressive cardiac dysfunction.

Epitranscriptomic modifications act as a critical layer of gene regulation to fine-tune essential biological processes (16). N6-methyladenosine (m⁶A) is the most abundant internal mRNA modification in eukaryotic cells and is deposited by the RNA methyltransferase 3 (METTL3)–METTL14 methyltransferase complex with WT1 associated protein (WTAP) (17, 18). This modification is interpreted by various m⁶A-binding “reader” proteins–such as YTH-domain family members–to regulate transcription (19–21), pre-mRNA processing (22, 23), mRNA stability (24–26), and translation (27–29). Conversely, m⁶A marks can be removed by demethylases, including fat-mass and obesity–associated protein (FTO) and AlkB homolog 5, RNA demethylase (ALKBH5) (30, 31). Through the dynamic progress of deposition, recognition and removal, m⁶A has been implicated in cancer, neurological disorders, and other human diseases (32–34).

Emerging evidence implicates m^6^A in regulating heart structure and function. Gene ontology (GO) analyses show that m⁶A modification in failing hearts is enriched on transcripts governing mitochondrial function and energy metabolism (35, 36), suggesting that altered m⁶A modification may contribute and impaired cardiac performance via altering mitochondrial function and metabolic process. Yet the clear evidence for this hypothesis remains unavailable. In addition, human and preclinical data reveal broad epitranscriptomic rewiring across heart failure phenotypes–including HFpEF– with m⁶A writer proteins including *Mettl3* and *Mettl4* upregulation (37, 38).

Consistent with these correlative observations, studies using genetically modified mouse models have directly established a causal role for m⁶A RNA modification in heart failure pathogenesis. Gain-of-function of METTL3 promotes pathological cardiac hypertrophy, whereas loss of METTL3 attenuates compensated hypertrophic growth and accelerates the transition to decompensated heart failure progress (35). Moreover, METTL14 overexpression suppresses exercise-induced physiological hypertrophy, while *Mettl14* knockdown via shRNA mitigates ischemia–reperfusion–induced cardiac injury and heart failure (39). Furthermore, cardiomyocyte-specific deletion of the m⁶A demethylase FTO prevents degradation of contractile gene transcripts; however, this paradoxically exacerbates cardiac dysfunction by impairing m⁶A-dependent translational efficiency of key cardiac proteins (36, 37). Together, these findings indicate that m⁶A serves as a dynamic regulatory mechanism that supports compensatory cardiac hypertrophy and safe guide cardiac homeostasis in physiological and pathological settings. However, the bona fide downstream targets of METTL14 that mediate its role in maintaining cardiac homeostasis remain largely unknown.

In this study, we have found that cardiomyocyte-specific ablation of *Mettl14* caused myocarditis and rapidly progressive dilated cardiomyopathy with early lethality, characterized by reduced left-ventricular ejection fraction (LVEF), left ventricular dilation, and posterior wall thinning. Conversely, *Mettl14* overexpression modestly enhanced systolic performance and LV wall thickness. Integrated MeRIP-seq and RNA-seq analyses revealed hypomethylated, upregulated transcripts enriched in the cGAS–STING–IFN-I axis in the *mettl14* knockout heart, establishing a direct link between METTL14-dependent m⁶A and IFN-I signaling pathways. Remarkably, interferon α/β Receptor subunit 1 (IFNAR1) deletion rescued the heart failure phenotype and prolonged survival in the in the *mettl14* background. Furthermore, METTL14 loss augmented receptor-interacting protein kinase 1 (RIPK1)-dependent necroptosis. Thus METTL14-mediated m⁶A modification restrains excessive innate immune activation and protects cardiomyocytes from RIPK1-mediated cell-death.

## Results

### Cardiomyocyte-specific deletion of *Mettl14* results in dilated cardiomyopathy with early lethality

To define how METTL14-mediated mRNA m^6^A regulated cardiac function, we generated a cardiomyocyte-specific *Mettl14* knockout (cmMettl14-/-) mouse line by crossing floxed *Mettl14* mice (40) with MLC2vKICre mice (41–43). MLC2vKICre efficiently deleted *Mettl14* in the heart without altering accumulation of WTAP or FTO proteins (Fig. 1A, B). METTL3 protein was modestly reduced, suggestive of destabilization of METTL3/METTL14 writer complex and resultant destruction of METTL3 in the *Mettl14* mutant. Dot-blot analysis of mRNA showed approximately 50% lower m^6^A levels of global mRNA in cmMettl14-/- vs the control hearts, validating that the tissue-specific knockout of *Mettl14* could effectively reduce the m^6^A level in the heart supporting the vigor of the subsequent physiological and mechanistic studies with the line are vigor (Fig. 1C).

**Figure 1:**
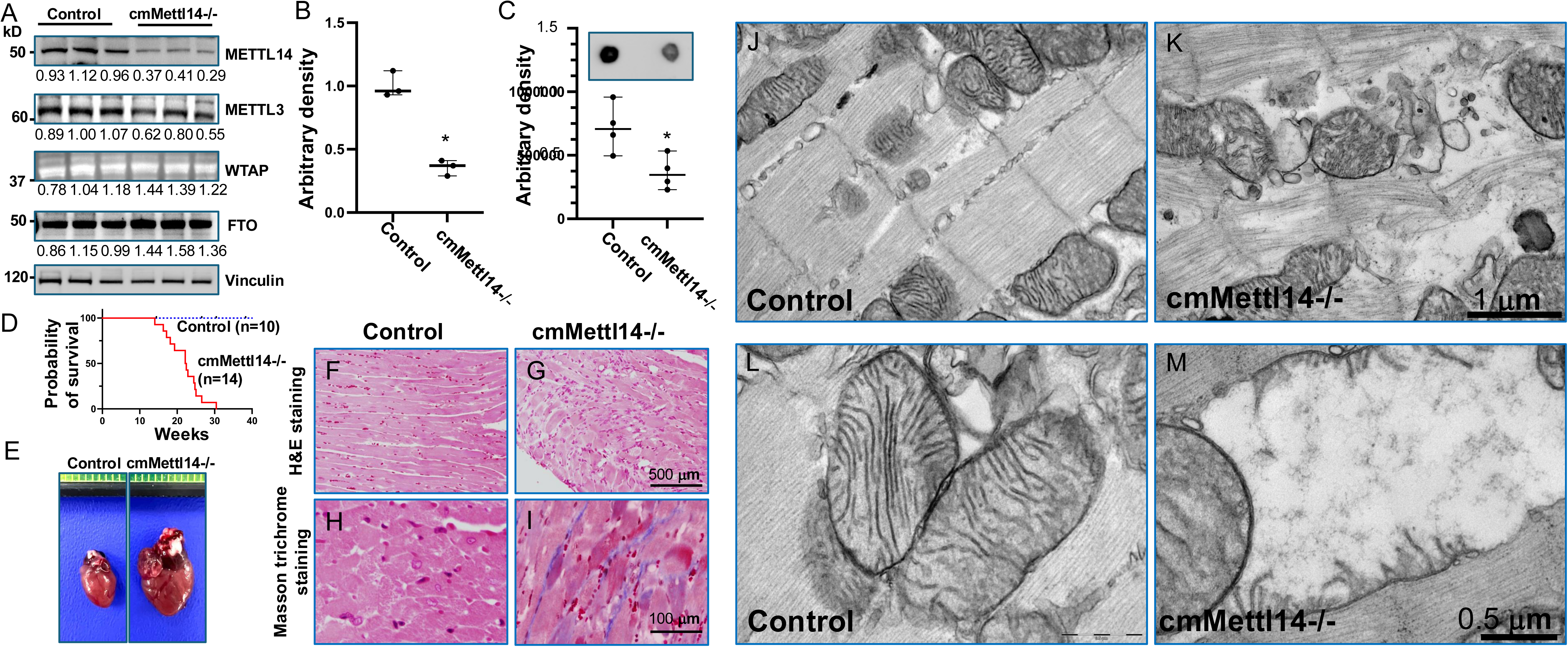
Cardiomyocyte-specific deletion of *Mettl14* causes severe dilated cardiomyopathy with early lethality. **(A, B)** Western blot analysis shows reduced METTL14 protein levels, while WTAP and FTO accumulation remained unchanged, in hearts from two-month-old knockout mice. (**B**) Quantification of METTL14 protein levels in control and knockout hearts. **(C)** Dot blot analysis and quantification show decreased global m^6^A modification in knockout hearts. **(D)** Kaplan–Meier survival curves demonstrate early mortality in *Mettl14* knockout mice. **(E)** Representative gross images of hearts from five-month-old control and knockout mice show cardiomegaly in *Mettl14* knockouts. **(F, G)** Hematoxylin and eosin (H&E) staining shows cardiomyocyte disorganized and inflammatory infiltration in *Mettl14* knockout hearts compared with control. **(H, I)** Masson trichrome staining shows interstitial fibrosis in *Mettl14* knockout hearts. (**J-M**) TEM images of control **(J, L)** and knockout hearts **(K, M)**. Control hearts show intact myofibrillar alignment and tightly packed mitochondrial cristae **(J)**, whereas knockout hearts exhibit loosely arranged myofibril **(K)**, swollen mitochondria, disrupted or absent cristae, and lucent mitochondrial matrix **(M)**. Data are shown as mean ± SD; ANOVA, *p* < 0.05, knockout vs. control.

We systemically accessed phenotypes of cardiomyocyte-specific *Mettl14* knockout mice. Survival analysis revealed early lethality beginning at ∼3 months, with 100% mortality by 7 months in cmMettl14-/- while the flox/flox control mice remain healthy (Fig. 1D). Gross examination of five-month-old survivors showed marked cardiomegaly for the mutant (Fig. 1E, right). Histology analysis demonstrated that cmMettl14-/- hearts displayed myocyte disarray with prominent inflammatory infiltrates, phenocopying the myocarditis, whereas the control hearts exhibited well-organized myocardium (Fig. 1F and G). Masson-trichrome staining clearly detected fibrosis in the cmMettl14-/- mice heart (Fig. 1I), but not in the control (Fig. 1H).

To determine effect of *Mettl14* deletion on the cardiomyocyte ultrastructure, we performed transmission electron microscopy (TEM) analysis. Control cardiomyocytes showed tightly packed, parallel myofibrils (Fig. 1J), while cmMettl14-/- hearts exhibited loosely arranged myofibrils, widened inter-myofibrillar spaces, and regions of myofibrillar dissolution with electron-lucent material (Fig. 1K). Mitochondria in controls contained intact, regularly spaced cristae (Fig. 1L); In contrast, they were swollen with disrupted/absent cristae and lucent, depleted matrices in cmMettl14-/- mice (Fig. 1M). Together, cardiomyocyte-specific loss of *Mettl14* reduced mRNA m^6^A modification and led to myocarditis, and severe dilated cardiomyopathy with early lethality.

### Inactivation of *Mettl14* in cardiomyocytes causes HF

To determine effect of METTL14 mediated m6A modification on heart function, we performed serial echocardiography in control and cmMettl14-/- mice at two, three, and four months of age. B-mode imaging revealed marked enlargment of left ventricle internal diameter (LVID) in cmMettl14-/- mice (Fig. 2B vs. 2A). Consistently, M-mode demonstrated increased LVID with reduced posterior wall thickness (PWT) in the cmMettl14-/- heart (Fig. 2D vs. 2C). Tissue-Doppler at the mitral valve annulus showed the decreased systolic (S) and early diastolic (e’) velocities, indicating impaired relaxation in the cmMettl14-/- hearts vs the controls (Fig. 2E, F). Quantitatively, LVEF was preserved at two months but declined sharply at three and four months in cmMettl14-/- mice (Fig. 2G), while LVIDd increased and PWTd decreased (Fig. 2H, I). The E/e′ ratio was elevated at three and four months (Fig. 2J), consistent with increased filling pressures. Together, these data show that the loss of METTL14 could cause progressive LV dilation with combined systolic and diastolic dysfunction, culminating in heart failure.

**Figure 2:**
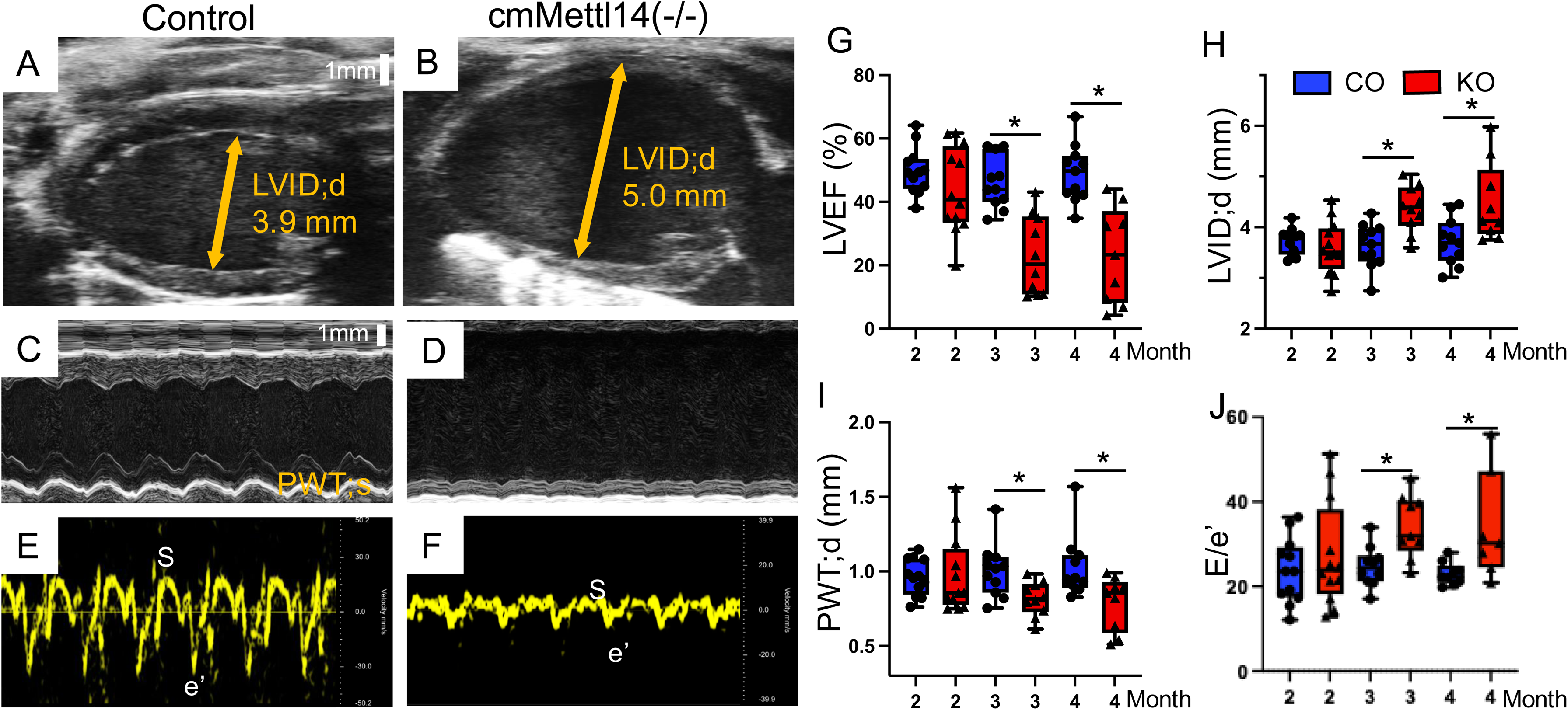
Cardiomyocyte-specific deletion of *Mettl14* induces severe HF. **(A, B)** Representative B-mode parasternal long axis and **(C, D)** M-mode parasternal short axis echocardiograms show LV dilation in knockout heart. **(E, F)** Representative tissue doppler traces at mitral valve annulus demonstrate severe diastolic dysfunction in knockout mice. **(G-J)** Quantification shows reduced LV ejection fraction (**G**), increased LV diameter (**H**), decreased LV posterior wall thickness during diastole (**I**), and elevated E/e′ ratio (**J**) in *Mettl14* knockout mice after three months. Data are shown as mean ± SD; ANOVA, *p* < 0.05, *Mettl14* knockout vs. control.

### Overexpression of *Mettl14* in cardiomyocytes induces cardiac hypertrophy

To define the gain-of-function effects of METTL14, we generated a cardiomyocyte-specific *Mettl14* overexpression (OE) mouse line (Cre-dependent transgene; Fig. 3A). Western blot assays showed that, METTL14 accumulation was increased by ∼50% in three-month-old OE mice versus controls (Fig. 3B, C). A reproducible, slower-migrating METTL14 band appeared in the OE hearts, suggestive of a potential but not yet identified posttranslational modification of METTL14 *in vivo.* An eGFP reporter confirmed cardiac-restricted expression–signal was present only in Cre-positive hearts and absent from skeletal muscle, lung, and other tissues (Fig. 3D). Echocardiography showed increased LVEF (Fig. 3E), reduced LVID in systole and diastole (Fig. 3F, G), and increased PWTs/d (Fig. 3H, I). Thus, in contrast to *Mettl14* loss, cardiomyocyte-specific overexpression of METTL14 could enhance contractile function and promote concentric remodeling consistent with cardiac hypertrophy.

**Figure 3:**
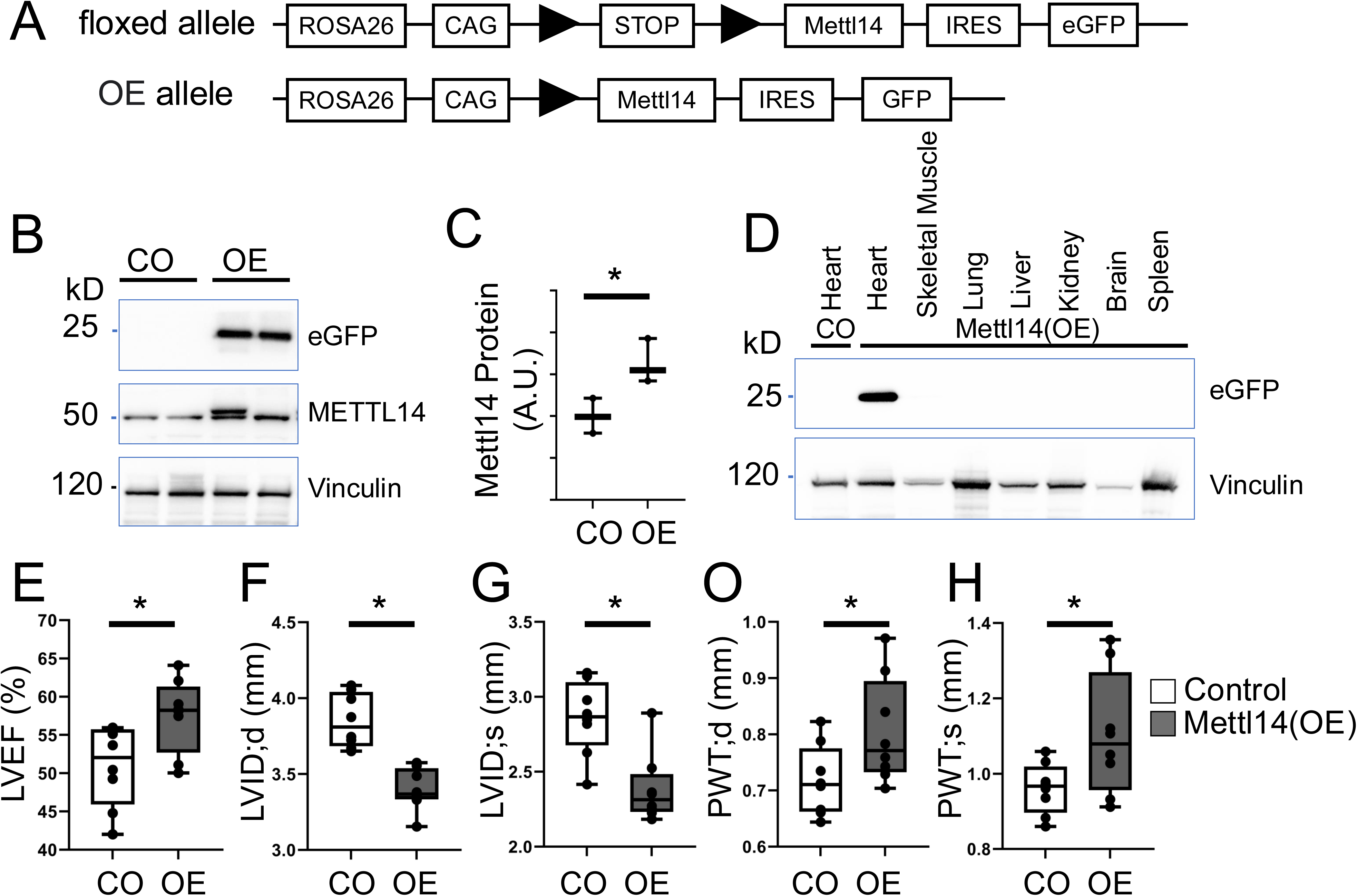
Overexpression of *Mettl14* in cardiomyocytes results in LV concentric hypertrophy. **(A)** Diagram illustrates the floxed control (CO) and *Mettl14* overexpression (OE) alleles. **(B-C)** Western blot analysis demonstrates increased METTL14 protein expression in hearts of three-month-old *Mettl14* OE mice compared to controls. **(D)** Exogenous METTL14 expression, detected via eGFP reporter, is specific to the heart and absent in other organs. **(E-H)** Echocardiographic analysis revealing enhanced left ventricular (LV) ejection fraction, reduced LV diameter, and increased LV posterior wall thickness during diastole and systole in *Mettl14* OE mouse hearts compared to controls. Statistical analysis: ANOVA, p < 0.05, OE vs. Control (CO).

### Loss of adaptive hypertrophic response to transvers aortic constriction (TAC) in cmMettl14-/- hearts

To test whether METTL14 was required for the adaptive hypertrophic response to pressure overload, we performed TAC in ten-week-old female mice. We chose female mice because they exhibited a milder baseline heart failure phenotype than males. We found that in two weeks after TAC challenge, the size of LVID was significantly increased in cmMettl14-/-hearts, compared to control hearts (Fig. 4A, B). The thickness of PWT was significantly decreased in cmMettl14-/- hearts (Fig. 4C, D). Quantitatively, LVEF was reduced (Fig. 4E), LVID in systolic and diastolic stage were increased (Fig. 4F, G), and PWTd/s were reduced (Fig. 4H, I) in cmMettl14-/- versus controls. Peak aortic velocity was comparably elevated in both groups, confirming similar TAC severity (Fig. 4J). In summary, the control hearts mounted the normal concentric hypertrophic remodeling seen, whereas the cmMettl14-/-hearts failed to thicken the LV wall and instead underwent dilation, leading to worsened systolic function. These findings indicate that the loss of *Mettl14* impairs adaptive hypertrophy and exacerbated TAC-induced heart failure.

**Figure 4:**
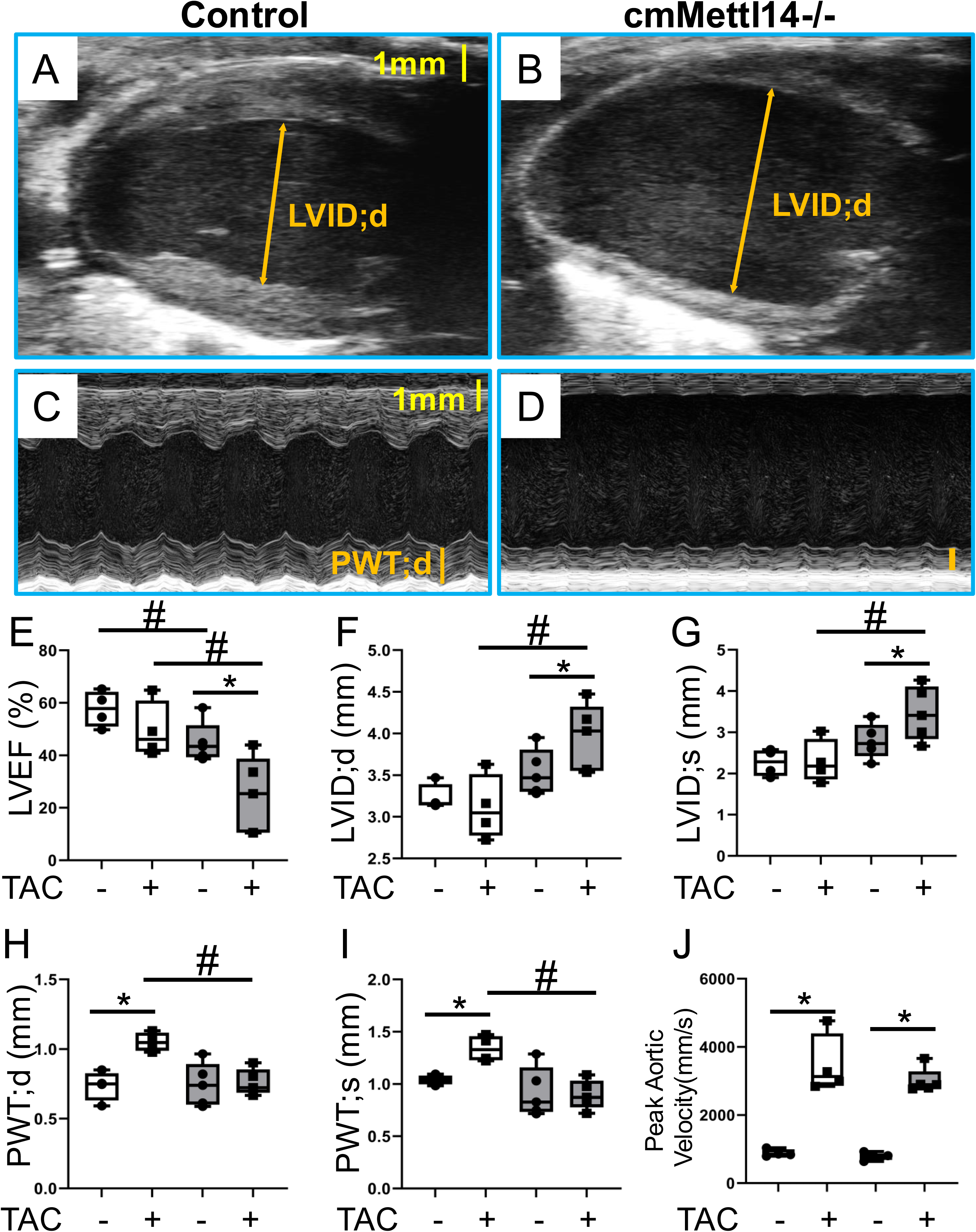
*Mettl14* deletion impairs adaptive cardiac remodeling in response to TAC-induced pressure overload in female mice. (A, B) Representative B-mode parasternal long-axis and **(C, D)** M-mode parasternal short-axis echocardiographic images of control **(A, C)** and cmMettl14-/-female mouse hearts **(B, D)** at two weeks post-TAC. **(E)** cmMettl14-/- mice displayed reduced LV ejection fraction two-week post-TAC, unlike control mice. **(F-I)** Control mice show increased LV posterior wall thickness and unchanged LV diameter, while cmMettl14-/- mice showed unchanged LV posterior wall thickness and increased LV diameter two-week post-TAC. **(J)** Both groups show elevated blood velocity in aortic arch two weeks post-TAC. ANOVA test: *p<0.05 after vs pre-TAC; #p<0.05 KO vs CO.

### Depletion of *Mettl14* in cardiomyocytes activates the cGAS-STING signaling pathway

We hypothesized that METTL14 deposited m^6^A on key transcripts to maintain the normal cardiac function, and that loss of METTL14 reduces this modification, leading to heart failure. To pinpoint the causative *bona fide* transcript targets of METTL14, we performed parallel RNA-seq and MeRIP-seq on eight-week-old cmMettl14-/- and control hearts, a time point prior to the onset of heart failure. RNA-seq analysis identified 805 differential expression genes (DEG), with 362 downregulated and 442 upregulated in cmMettl14-/- hearts compared to control heart (Fig. 5A). The GO analysis linked the downregulated genes to the biological events of heart contraction and tRNA processing (Fig. 5B). In contrast, the most significant upregulated genes were predominantly implicated in inflammatory (Fig. 5C). Specifically, 7 of the top 10 enriched pathways were related to inflammatory activation, including regulation of immune effector functions, cytokine production, and leukocyte migration in cmMettl14-/- hearts. The aberrant inflammatory activation revealed by RNA-seq is consistent with our histological analysis, which demonstrated myocarditis in cmMettl14-/- hearts (Fig. 1G). Together, these findings indicate that METTL14 functions as a key regulator that restrains innate immune responses to maintain cardiac homeostasis.

**Figure 5:**
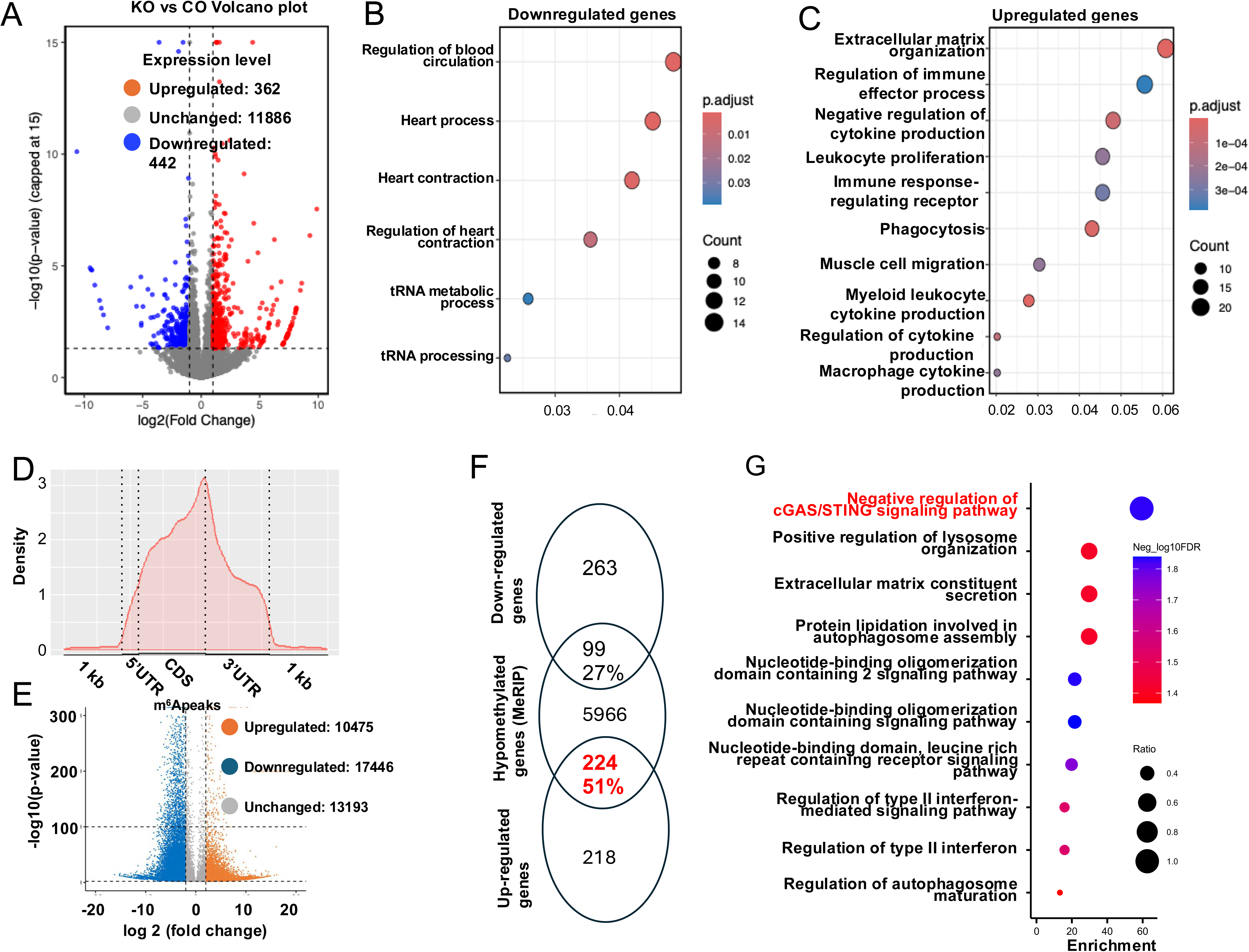
Characterization of m^6^A modified transcripts. **(A)** RNA-seq volcano plot identify 804 DEGs: 362 downregulated and 442 upregulated transcripts in cmMettl14-/- vs control hearts. **(B, C)** GO analysis reveals that downregulated genes are enriched for cardiac contraction/tRNA processing, whereas upregulated genes are enriched for immune effector processes and cytokine production. **(D)** Distribution of m^6^A peaks across the length of mRNA transcripts in the control mouse hearts. **(E)** MeRIP-seq volcano plots demonstrate widespread loss of m^6^A peaks in cmMettl14-/- hearts. **(F)** Overlap analysis of DEGs with hypomethylated transcripts reveals a significant intersection, including 224 hypomethylated/upregulated genes. **(G)** GO/pathway analysis of these hypomethylated-and-upregulated genes highlights cGAS–STING signaling among the top enriched categories.

Our parallel MeRIP-seq results presented that m^6^A peaks were mostly distributed on 3’ untranslated regions and stop codons’ segments (Fig. 5D). This result was in lines with previous reports (36), justifying the reliability of the MeRIP-seq. Importantly, we detected 17,446 hypomethylated peaks across 6,279 mRNAs (≥2-fold, p<0.01 upon *Mettl14* inactivation). We also called apparent “hypermethylated” peaks (n=10,475) in the cmMettl14-/- hearts. This scenario was likely artifacts from the computational algorithm that utilized internal normalization to calculate the relative alternation the m^6^A level of transcripts inside the samples (Fig. 5E) and thus were not pursued. Venn graphing between DEGs from RNA-seq and hypomethylated transcripts from MeRIP-seq revealed that 224 of 442 upregulated genes (51%) were hypomethylated (hypergeometric test p=6.964e-13<0.05), whereas only 99 of 362 downregulated genes (27%) were hypomethylated (p=0.998) (Fig. 5F). Thus, the transcripts that increase in abundance were obviously associated with the reduced m^6^A modification, suggesting that m^6^A loss would stabilize and/or accumulate the harbored mRNAs. This result mirrored the earlier reports that show m^6^A is a negative regulator for mRNA accumulation (24, 44, 45).

Importantly, GO enrichment analysis of the 224 hypomethylated and upregulated genes revealed a strong overrepresentation of a pathway associated with innate immune activation (Fig. 5G). Remarkably, the top enriched pathway was classified as the negative regulation of the cGAS-STING signaling pathway. Although this may seem counterintuitive at first glance, such enrichment often reflects compensatory transcriptional responses triggered by upstream overactivation. Consistent with this idea, we observed that cardiomyocyte-specific loss of METTL14 leads to mitochondrial damage and myocarditis (Fig. 1G, M), suggesting activation of innate immune signaling mediated by the cGAS-STING pathway in cmMettl14-/- hearts. Supporting this interpretation, additional enriched pathways included type II interferon signaling (IFN-γ pathway) and lysosome maturation, both of which are closely linked to cytosolic DNA sensing and downstream innate immune responses (9).

Among the genes that contributed to the cGAS-STING–related enrichment, Irgm*1*, *Irgm2*, and *Irgm3* displayed most significant upregulation. IRGM1 is a key regulator of mitochondrial integrity and mitophagy, and its precise expression level is critical for maintaining mitochondrial membrane potential and overall mitochondrial homeostasis. Excessive IRGM1 expression has been shown to impair mitochondrial quality control, thereby facilitating mtDNA leakage into the cytosol, a potent activator of the cGAS-STING pathway (46–48). The concurrent enrichment of positive regulation of lysosome maturation further supports increased mitophagy-related stress responses in METTL14-deficient cardiomyocytes.

Importantly, the GO terms also indicated significant upregulation of the IFN-II signaling pathway. Given that IFN-I and IFN-II converge on shared downstream effectors such as STAT1 (9), the elevation of IFN-II–related pathway strongly suggests a broader interferon-primed state. Together with the transcriptional signatures of mitochondrial stress and cGAS-STING activation, these data support a model in which loss of METTL14 disrupts mitochondrial homeostasis by increasing IRGM1 expression, promotes mtDNA release, activates the cGAS-STING pathway, and consequently drives IFN-I upregulation in the myocardium.

### cGAS-STING pathway activation and increased IFN-I in cmMettl14-/- hearts

Activation of the cGAS-STING pathway is a central mechanism driving IFN-I upregulation. Here in our hands, GO analysis highlighted the changes of cGAS-STING pathway in cmMettl14-/- hearts. These results prompted us to further assess expression of interferon stimulated genes (ISG), the responses of cGAS-STING activation and IFN-I upregulation, in our RNA-seq database. Indeed, the depletion of *Mettl14* significantly induced expression of a wide array of ISGs (Fig. 6A), indicative of aberrant cGAS-STING-IFN-I activation in cmMettl14-/-hearts. These results were further validated by RT-qPCR assays that showed elevation of IFN-I transcripts and ISGs in whole hearts (Fig. 6B) and in isolated cardiomyocytes (Fig. 6C). In lines with the transcript upregulation, the accumulation of STING and IRF3 proteins were observed in cmMettl14-/- hearts (Fig. 6D). Together, these data indicate that loss of METTL14 activates the cGAS-STING axis in cardiomyocytes, drives IFN-I production, and likely contributes to IFN-I-mediated heart failure.

**Figure 6:**
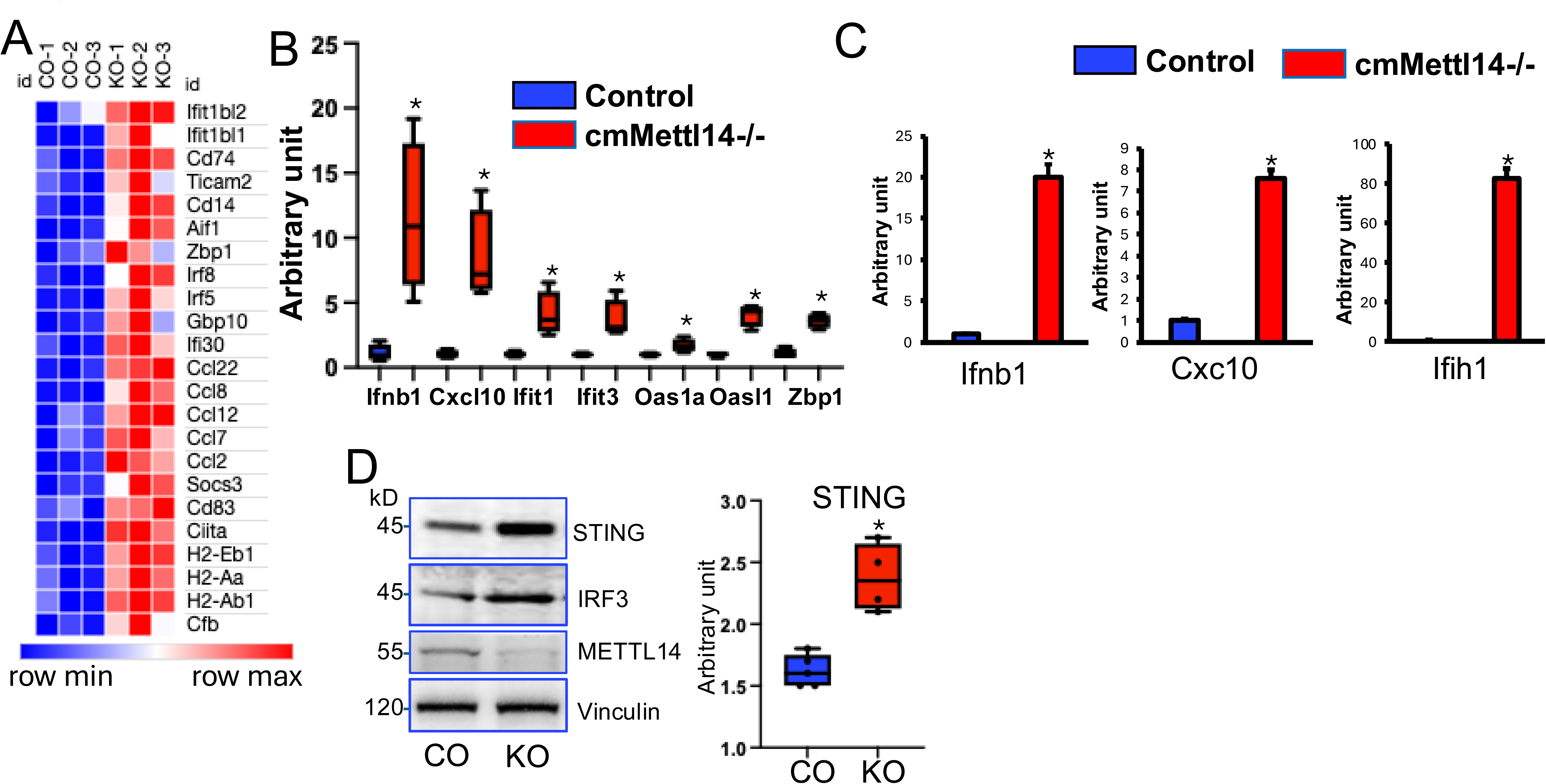
Inactivation of *Mettl14* induced STING, *Ifnb1* and *ISG* upregulation. **(A)** Heatmap analysis of ISG expression from RNA-seq data in cmMettl14-/- (KO) versus control (CO) hearts. **(B, C)** RT-qPCR analysis show upregulated *ISG* gene expression in cmMettl14-/- hearts (C) and isolated cardiomyocytes (D) compared to controls. **(D)** Western blot analysis demonstrates elevated STING and IRF3 protein levels in cmMettl14-/- (KO) hearts compared to controls (CO). * p<0.05 KO vs CO.

### Inactivation of *Ifnar1* rescued *Mettl14* deletion induces heart failure

To investigate whether the aberrant activation of IFN-I derives for heart failure in *Mettl14* deletion mice, we crossed cmMettl14-/- mice with IFN-I receptor I (IFNAR1) knockout mice. We found that deletion of *Ifnar1* significantly extended survival rates of mice in the cmMettl14-/- background compared to both the parental lines and the sibling heterozygous controls (Fig. 7A). For detail, more than 75% *Ifnar1* and *Mettl14* double knockout mice were alive, whereas all cmMettl14-/- mice died within 33 weeks. Furthermore, we conducted echocardiography assays and found significant increase in LVEF (Fig. 7B) but a decrease in LVID in the *Ifnar1* and *Mettl14* double knockout mice (Fig. 7C, D), compared to cmMettl14-/-. These results indicated that inactivation of *Ifnar1* could indeed rescue the heart failure phenotype in the *Mettl14*-depletion background, with a further indication that METTL14 controls heart homeostasis via restraining IFN-I signaling.

**Figure 7.**
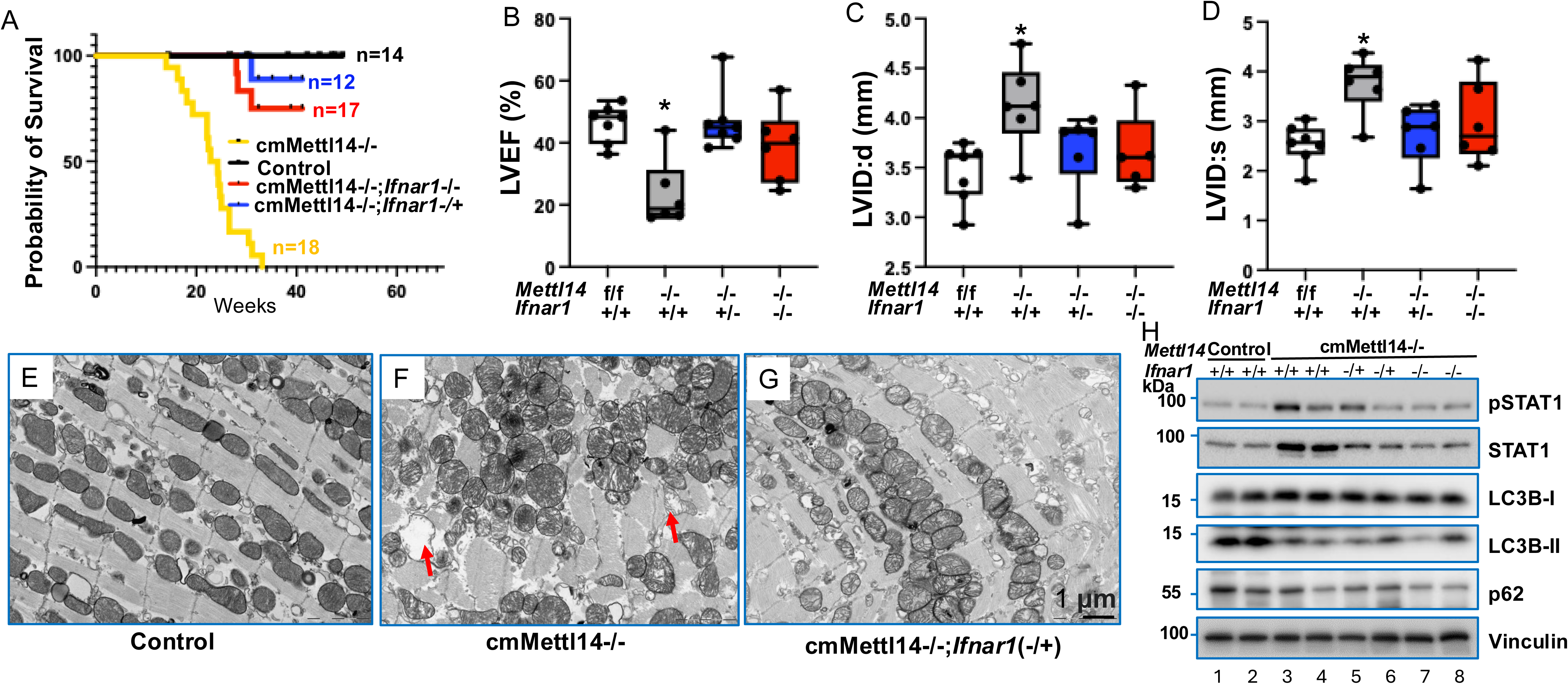
Inactivation of *Ifnar1* rescues *Mettl14* deletion induced HF. (**A**) Kaplan–Meier curves show that *Ifnar1* knockout rescues the early lethality of cmMettl14-/- mice. (**B-D**) Echocardiography presents that LVEF is increased (B) and LVIDd/s are decreased (C, D) in *Ifnar1; Mettl14* double-knockout mice compared with *cmMettl14-/-*. (**E-G**) TEM images of control (**E**), cmMettl14-/- (**F**), and *Ifnar1; Mettl14* double knockout (**G**) hearts; red arrows highlight the swollen mitochondria. (**H**) Immunoblot of autophagy markers LC3B and p62 in control, cmMettl14-/-, and *Ifnar1*; *Mettl14* double knockout hearts. *Ifnar1+/+*: wild type; *Ifnar1-/+*: heterozygous; *Ifnar1*-/-: knockout. ANOVA test: *: cmMettl14-/- vs others, p<0.05.

To further validate the protective effect of *Ifnar1* deletion on *Mettl14*-deficiency–induced cardiomyocyte damage, we performed transmission electron microscopy (TEM) to assess ultrastructural changes. TEM analysis of cmMettl14-/- hearts revealed pronounced mitochondrial swelling, cristae disruption, and disorganized myofibrils (Fig. 7F), abnormalities that were absent in control hearts (Fig. 7E). Strikingly, these ultrastructural defects were substantially ameliorated in *Ifnar1/Mettl14* double-knockout hearts (Fig. 7G), with restoration of mitochondrial morphology and preservation of myofibril organization. These findings demonstrate that loss of Ifnar1 markedly mitigates the ultrastructural damage caused by Mettl14 deletion, further supporting the central role of IFN-I signaling in mediating Mettl14-deficiency–induced cardiomyocyte injury.

Given that METTL14 deletion activates IFN-I signaling, we next examined alterations in downstream IFN-I signal transduction that is critical for METTL14-mediated regulation of heart failure pathogenesis. Because signal transducer and activator of transcription 1 (STAT1) is the principal effector of IFNAR1 and a key mediator of interferon-stimulated gene expression, we assessed STAT1 activation in our models. Cardiomyocyte-specific deletion of *Mettl14* led to a marked increase in both total and phosphorylated STAT1 protein levels, consistent with enhanced IFN-I pathway activation (Fig. 7H, lanes 3–4 vs. 1–2). Notably, genetic ablation of *Ifnar1* completely eliminated this STAT1 upregulation (Fig. 7H, lanes 5–8 vs. 3–4). These findings demonstrate that METTL14 loss augments IFN-I production and IFNAR1-dependent signaling, thereby contributing to the progression of heart failure (Fig. 7H, top two panels). The inflammatory phenotype observed in METTL14-deficient hearts mirrors the immunopathology of ICI-induced myocarditis, where IFN-γ–producing T cells and macrophages drive cardiotoxicity (49, 50). Although IFN-I and IFN-γ signal through different receptors, both converge on STAT1 to amplify ISG responses and cytotoxic effector mechanisms. The complete protection afforded by *Ifnar1* deletion in our model demonstrates that cardiomyocyte-intrinsic IFNAR1–STAT1 signaling is a major driver of myocarditis, even in the absence of exogenous immune stimulation. These results reveal METTL14 as a previously unrecognized molecular brake that restrains intrinsic antiviral signaling and maintains cardiac immune tolerance.

In addition to delineating alterations in the cGAS-STING-IFN-I axis, GO analysis also revealed significant enrichment of pathways related to autophagy and lysosomal maturation (Fig. 5G). Because efficient autophagy/mitophagy is essential for clearing dysfunctional mitochondria and thereby preventing aberrant activation of the cGAS-STING pathway (48), we next assessed whether *Mettl14* deletion disrupts autophagic flux. LC3B and p62 were selected as canonical molecular markers for monitoring autophagy: LC3B lipidation (conversion of LC3B-I to LC3B-II) reflects autophagosome formation, whereas p62 accumulation or depletion indicates autophagosome cargo processing and lysosomal degradation efficiency (51, 52). Cardiomyocyte-specific inactivation of Mettl14 markedly reduced LC3B lipidation, as shown by diminished conversion of LC3B-I to LC3B-II (Fig. 7H, third and fourth panels). In parallel, p62 protein levels were substantially decreased in cmMettl14-/- hearts (Fig. 7H, fifth panel), together demonstrating impaired autophagic/mitophagic flux. Notably, loss of IFNAR1 did not restore these defects, indicating that disrupted autophagy/mitophagy lies upstream of IFN-I production and functions as an initiator, rather than a downstream effector, of IFN-I pathway activation.

### Cardiomyocyte-specific *Mettl14* deletion induces necroptosis

Inflammation-driven cell death is a key driver of heart failure, promoting cardiomyocyte loss and amplifying sterile inflammation in a vicious cycle of injury and maladaptive remodeling (5, 53). Since inactivation of *Mettl14* resulted in myocarditis and dilated cardiomyopathy accompanied by elevated IFN-I production, we next investigated whether METTL14 deficiency directly drives cardiomyocyte death and/or acts through additional, previously unrecognized genetic pathways. To address this, we performed Western blot analyses of key markers of apoptosis, pyroptosis, and necroptosis. *Mettl14* deletion markedly reduced poly (ADP-ribose) polymerase (PARP) cleavage, gasdermin D cleavage, and IL-1β accumulation (Fig. 8A), indicating suppression of both apoptotic and pyroptotic pathways in METTL14-deficient hearts.

**Figure 8.**
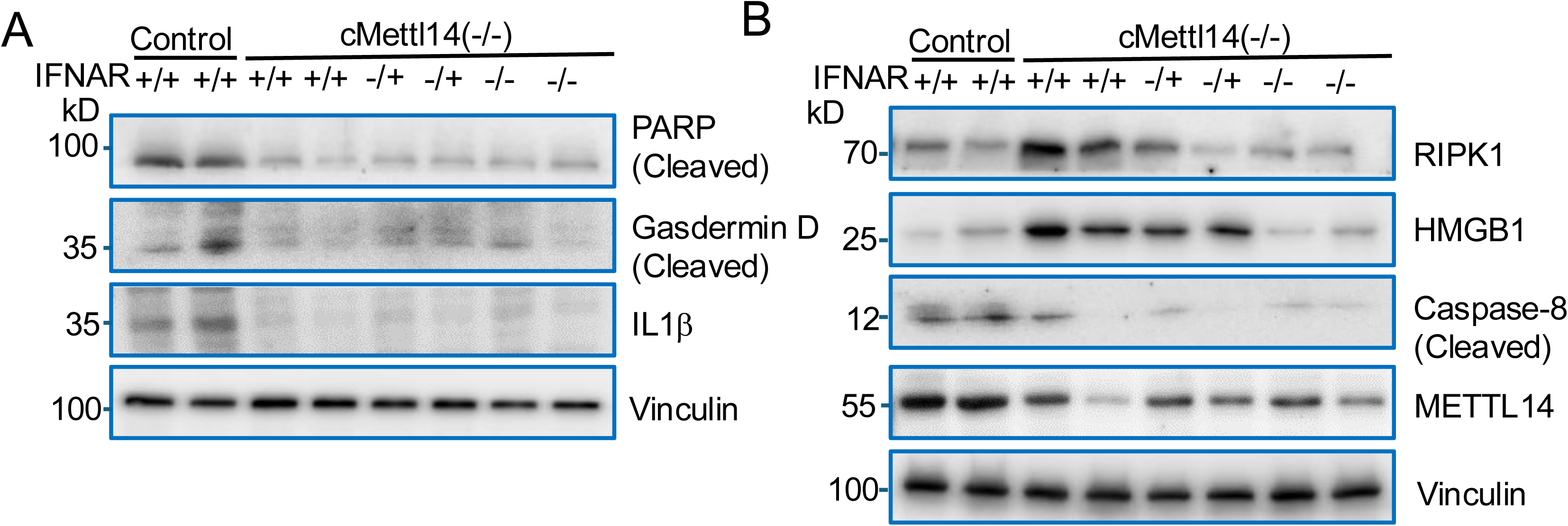
METTL14 deficiency promotes necroptosis in the heart. (**A**) Immunoblotting shows that *Mettl14* inactivation reduces cleavage of PARP, gasdermin D, and IL-1β, indicating dampened apoptosis/pyroptosis. (**B**) *Mettl14* deletion increases RIPK1, HMGB1, and Caspase-8, consistent with activation of necroptotic signaling.

Because necroptosis has emerged as a major form of inflammatory cell death in the heart, and RIPK1–RIPK3–MLKL signaling governs its initiation, we specifically examined RIPK1 as a central upstream regulator of necroptosis. In parallel, we assessed high-mobility group box 1 (HMGB1), a prototypical alarmin that is passively released during necroptotic but not apoptotic cell death and thus serves as a functional readout of necroptosis-driven damage-associated molecular pattern (DAMP) release. METTL14 loss resulted in upregulation of RIPK1 and HMGB1 (Fig. 8B), providing strong evidence for enhanced necroptotic signaling and inflammatory DAMP release. Moreover, RIPK1 was among the 224 hypomethylated and upregulated transcripts identified in cmMettl14-/- hearts (Fig. 5F), suggesting that loss of METTL14-mediated m⁶A modification stabilizes RIPK1 mRNA, leading to increased RIPK1 protein accumulation and activation of necroptotic signaling.

These molecular signatures also align with our GO enrichment analysis showing the elevation of nucleotide-binding oligomerization domain containing signaling pathway (NOD1)/NOD2, which in turn heightens intracellular danger surveillance and amplifies pro-inflammatory signaling, thereby creating a necroptosis-permissive environment in *Mettl14*-deficient hearts. Notably, genetic inactivation of *Ifnar1* rescued the upregulation of RIPK1 and HMGB1, indicating that IFNAR1-dependent signaling specifically amplifies RIPK1-mediated necroptosis rather than globally regulating cell death pathway components.

Caspase-8 is a master molecular switch that governs the balance among apoptosis, pyroptosis, and necroptosis (54–56). By activating executioner caspases-3 and -7, caspase-8 promotes apoptotic cell death, while simultaneously restraining necroptosis through proteolytic cleavage of RIPK1 and RIPK3 (54, 56, 57). Loss or inhibition of caspase-8 therefore disables apoptotic pathways and unleashes RIPK1-dependent necroptotic signaling, positioning caspase-8 as a pivotal regulator of cell fate and inflammation in the failing heart (56). In our study, cardiomyocyte-specific inactivation of *Mettl14* significantly reduced caspase-8 protein levels, and this defect was not restored by *Ifnar1* deletion. These findings strongly support a model in which METTL14 deficiency decreases caspase-8, thereby suppressing apoptosis and pyroptosis while promoting RIPK1-mediated necroptosis. This shift in the cell-death landscape likely contributes to sterile inflammation through the release of HMGB1 and other damage-associated molecular patterns. The rescue of necroptosis by *Ifnar1* deletion further underscores a feed-forward loop linking IFN-I signaling, RIPK1 activation, and inflammatory injury.

Take together, our findings demonstrate that cardiomyocyte-specific METTL14 deficiency disrupts mitochondrial homeostasis, activates the cGAS–STING–IFN-I axis, and promotes a necroptosis-dominant inflammatory environment that further drives myocarditis and dilated cardiomyopathy. Mechanistically, METTL14 loss impairs autophagy/mitophagy, enhances NOD1/NOD2-mediated danger sensing, and activates RIPK1-dependent necroptosis, accompanied by increased HMGB1 release. Importantly, genetic inactivation of *Ifnar1* rescues IFN-I signaling and necroptosis, establishing IFNAR1-dependent innate immune activation as a central mechanism underlying METTL14 deletion–induced heart failure (Fig. 9).

**Figure 9.**
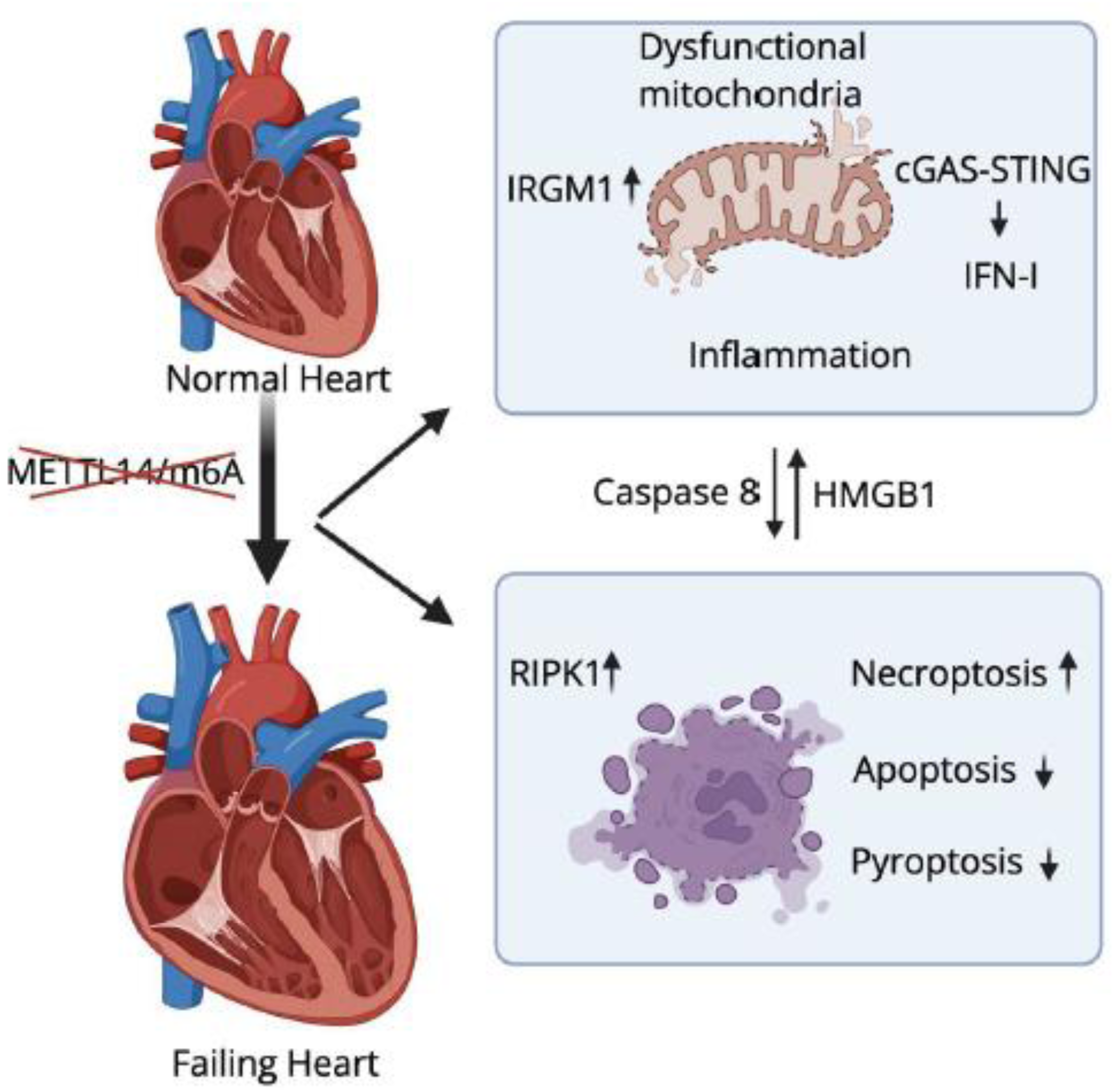
Proposed model of METTL14-mediated m6A modification in heart failure. Loss of METTL14 alters m6A modification, leading to mitochondrial dysfunction, cGAS-STING-IFN-I activation, and necroptosis. These parallel pathways converge to drive cardiomyopathy and heart failure.

## Discussion

RNA m⁶A methylation has emerged as a regulatory factor for diverse physiological and disease contexts.(17, 18, 58). Here, we identify METTL14-dependent m⁶A modification as an essential intrinsic checkpoint that maintains cardiomyocyte immune quiescence, mitochondrial homeostasis, and cell survival. Cardiomyocyte-specific deletion of *Mettl14* triggered spontaneous IFN-I activation, severe myocarditis, progressive dilated cardiomyopathy, and early mortality. Strikingly, genetic ablation of *Ifnar1* rescued the lethal phenotype, restoring cardiac mitochondrial and sarcomere structure and function while suppressing necroptosis (Fig. 9). These findings reveal a previously unrecognized role of METTL14 in restraining pathological IFN-I activation and establishing a direct mechanistic link between RNA epitranscriptomic dysregulation, myocarditis, dilated cardiomyopathy, and heart failure.

### m⁶A remodeling in heart failure and therapeutic implications

Previous studies have reported increased m⁶A and METTL14 levels in failing human, mouse, and pig hearts, suggesting active m⁶A remodeling during disease progression (35). Paradoxically, both increased and decreased m⁶A levels result in heart failure: METTL3/METTL14 loss is detrimental (35), yet FTO loss, which increases m⁶A, is also harmful (36, 37). These findings indicate that m⁶A exerts both protective and pathogenic effects depending on the target transcripts, emphasizing the need to identify context-specific m⁶A-dependent regulators that determine cardiomyocyte fate.

Here parallel RNA-seq and MeRIP-seq analyses pinpointed the authentic m^6^A targeted mRNA in the heart. Consistent with previous reports (36), we observed that the number of altered m^6^A peaks substantially exceeded the number of transcripts with changed expression levels. This contrast supports that m^6^A modification acts beyond the regulation of transcript level exemplified by mRNA translation–during heart failure progression. Although prior MeRIP-seq studies have been performed in healthy heart and human end-stage heart failure samples, similar analyses have not been conducted in genetic knockout mice. Our parallel *in vivo* RNA-seq and MeRIP-seq analyses in m^6^A modification altered hearts provided comprehensive data regarding the true m^6^A targets. Our data presented that more 51% upregulated mRNAs are hypomethylated in cmMettl14-/- heart, thereby providing direct evidence linking m^6^A modification to RNA stability regulation in cardiac pathophysiology.

### METTL14–m⁶A preserves mitochondrial–immune homeostasis

Interferons act as a double-edged sword in the heart: beneficial during acute viral infection but detrimental when chronically elevated (9). Sustained IFN-I and IFN-γ signaling, as observed in viral myocarditis, ICI–associated myocarditis, and non-ischemic cardiomyopathy, drives maladaptive remodeling, protein synthesis suppression, and cardiomyocyte loss (9, 49, 50, 59, 60). Consistent with this paradigm, *Mettl14*-deficient hearts exhibited strong upregulation of IFN-stimulated genes (ISGs) and heightened IRF3 and STING expression, whereas inactivation of *Ifnar1* rescued Mettl14 deletion induced heart failure, indicating robust activation of cGAS-STING-IFN-I pathways.

Our data collectively support a model in which METTL14 suppresses aberrant IFN activation through broader control of mitochondrial homeostasis. METTL14 deficiency caused striking mitochondrial swelling, cristae disruption-all hallmarks of organelle stress. In parallel, autophagy/mitophagy flux was impaired, as evidenced by reduced LC3B cleavage and altered p62 accumulation. Because both impaired mitochondria and interfered mitophagy process contribute the accumulation of mtDNA in the cytosol and led to cGAS–STING pathway activation. Indeed, *Ifnar1* deletion rescued inflammatory signaling and necroptosis but did not correct the underlying mitophagy impairment, placing mitochondrial dysfunction upstream of IFN-I induction. Together, these findings delineate a METTL14–m⁶A–mitochondria axis that maintains cytosolic DNA homeostasis and prevents inappropriate innate immune activation in the heart.

### The cross talk between the activated IFN-I and RIPK1 induced necroptosis

One major novel finding of our study is that cardiomyocyte-specific deletion of *Mettl14* induces aberrant type I interferon (IFN-I) activation and RIPK1-mediated necroptosis, which together constitute a central pathogenic axis driving heart failure. Loss of METTL14-dependent m⁶A modification results in inappropriate activation of innate immune signaling, characterized by robust IFN-I production and IFNAR1-dependent downstream signaling. IFN-I signaling is well established to transcriptionally upregulate key components of the necroptotic machinery, including RIPK1, RIPK3, and MLKL, thereby sensitizing cardiomyocytes to programmed necrotic cell death under inflammatory stress. In the absence of METTL14, RIPK1 is both hypomethylated and upregulated, indicating that impaired m⁶A-dependent RNA turnover directly contributes to RIPK1 accumulation at both the transcript and protein levels. Elevated RIPK1 not only promotes necrosome assembly but also amplifies inflammatory signaling, establishing a feed-forward loop between IFN-I activation and necroptosis. Importantly, genetic ablation of Ifnar1 markedly attenuates RIPK1 upregulation, suppresses necroptotic signaling, and rescues cardiac dysfunction and survival, firmly positioning IFN-I signaling upstream of RIPK1-mediated necroptosis in this model. Collectively, these findings identify METTL14 as a critical epitranscriptomic brake that restrains IFN-I–RIPK1 crosstalk, thereby protecting cardiomyocytes from inflammatory necroptotic death and limiting heart failure progression. Although *Ifnb1* has been reported as an m⁶A-regulated transcript in other biological contexts (61, 62), it was not hypomethylated in our MeRIP-seq dataset, suggesting that its upregulation in cmMettl14-/-hearts may not reflect a direct m⁶A-dependent mechanism. Instead, METTL14 loss–induced mitochondrial dysfunction and impaired autophagy/mitophagy flux likely promote cytosolic mtDNA accumulation and subsequent activation of the cGAS–STING pathway, thereby driving secondary IFN-I induction.

### METTL14-dependent m⁶A supports adaptive hypertrophy and protects against heart failure

Our data further reveal that METTL14 is required for compensated hypertrophic remodeling. Loss of METTL14 significantly suppressed TAC-induced hypertrophy yet accelerated heart failure progression. RNA sequencing revealed broad downregulation of mitochondrial metabolic and structural genes, consistent with impaired mitochondrial capacity to support the increased energetic and biosynthetic demands of hypertrophy. Together, these results indicate that METTL14 orchestrates both metabolic adaptation and immune restraint during cardiac stress.

Interestingly, although both METTL3 and METTL14 deficiency led to eccentric hypertrophy and heart failure, the phenotype of Mettl14-deficient mice is more severe. This may reflect differences in recombination efficiency or genetic background, but could also suggest that METTL14 performs nonredundant structural or catalytic functions within the m⁶A writer complex. Supporting this, METTL14 loss reduces global m⁶A levels more dramatically than METTL3 loss in cultured cells (63), implying a more indispensable role for METTL14 in guiding m⁶A deposition.

In conclusion, our study provides a framework for such investigations by integrating transcriptome-wide m⁶A mapping, mitochondrial assays, and immunometabolic analysis. Collectively, our findings establish METTL14 as a master regulator of cardiomyocyte-intrinsic immunity, mitochondrial homeostasis, and cell-death decision-making. They further suggest that therapeutic modulation of m⁶A, or targeted inhibition of IFN-I or necroptotic signaling, may hold promise for treating inflammatory cardiomyopathies, including ICI-induced myocarditis and viral or autoimmune myocarditis.

## Material and Methods

### Mice

Cardiomyocyte-specific *Mettl14* knockout (cmMettl14-/-) mice were generated by crossing Mettl14^fl/fl^ mice (40) with MLC2vKICre transgenic mice (41, 42, 64). To assess the role of IFN-I signaling, cmMettl14-/- mice were crossed with *Ifnar1*-/- mice to generate double-knockout mice. All mice were maintained on a C57BL/6J and 129 mixture background and housed in a pathogen-free vivarium under a 12-hour light/dark cycle with ad libitum access to food and water. Both male and female mice were used. All animal procedures were approved by the Texas A&M University Institutional Animal Care and Use Committee (IACUC).

### Genotyping

Genomic DNA was isolated from ear or tail biopsies using standard hot sodium hydroxide and Tris method (65). PCR was performed using primers specific for *Mettl14* floxed alleles **(5’-**ACTTCACTTCCAACGCAGGT**-3’, 5’-**ACATGCAGAGTCAGCACCT**-3’)**, *Ifnar1* floxed alleles (Common: 5’-CGAGGCGAAGTGGTTAAAAG-3’, Wild type reverse: 5’-ACGGATCAACCTCATTCCAC-3’, Mutant: 5’-AATTCGCCAATGACAAGACG-3’) and Cre allele (5’**-**CGCAGAACCTGAAGATGTTCGCGATTA-3’, 5’-TCTCCCACCGTCAGTAC GTGAGATATC-3’). PCR products were resolved using agarose gel electrophoresis.

### Transverse aortic constriction (TAC) in mice

All animal experiments were conducted in accordance with the *Guide for the Care and Use of Laboratory Animals* (NIH Publication No. 85-23, revised 1996). Transverse aortic constriction (TAC) surgery was performed using a minimally invasive approach. Briefly, 10-week-old female mice were anesthetized by inhalation of 1–1.5% isoflurane. A 27-gauge needle was positioned adjacent to the transverse aorta between the right innominate and left common carotid arteries, and a 6-0 non-absorbable suture was tied securely around both the aorta and the needle; the needle was then removed to generate a standardized constriction. Sham-operated mice underwent an identical surgical procedure without aortic banding.

### Echocardiography

Transthoracic echocardiography was performed using a Vevo3100 imaging system equipped with an MX550D probe (VisualSonics). Mice were lightly anesthetized with 1–1.5% isoflurane to maintain heart rates between 450–550 bpm. M-mode images were acquired at the mid-papillary level to measure left ventricular (LV) posterior wall thickness, LV internal diameter (LVID), and ejection fraction (EF) using VevoLAB software.

### RNA isolation and mRNA purification

Total RNA was isolated from left ventricular tissue using TRI Reagent™ (Thermo Fisher Scientific) according to the manufacturer’s instructions. Poly(A)⁺ mRNA was subsequently purified from total RNA using oligo (dT) magnetic beads (Thermo Fisher Scientific) following the manufacturer’s protocol.

### m^6^A blot analysis

RNA samples were quantified by UV spectrophotometry and denatured at 95 °C for 3 min. The m⁶A dot blot assay was performed as previously described (66). Briefly, purified mRNA samples were immobilized onto Hybond-N⁺ membranes (GE Healthcare, USA) by UV crosslinking. Membranes were then blocked with 5% nonfat dry milk in 1× PBST for 1 h at room temperature and incubated overnight at 4 °C with an anti-m⁶A antibody (1:5,000; Cell Signaling Technology, Cat. #56593). After washing, membranes were incubated with HRP-conjugated anti-rabbit IgG secondary antibody (Cell Signaling Technology) for 1 h at room temperature. Signals were detected using SuperSignal™ West Pico PLUS Chemiluminescent Substrate (Thermo Fisher Scientific, Waltham, MA) and visualized with a Bio-Rad imaging system.

### Histology and Immunohistochemistryanalysis

Hearts were perfused with PBS, fixed in 4% paraformaldehyde overnight, dehydrated, and embedded in paraffin. Sections (4 μm) were stained with hematoxylin and eosin (H&E) or Masson’s trichrome to evaluate myocarditis severity and fibrosis.

### Transmission Electron Microscopy (TEM)

Small pieces of LV myocardium (1 mm³) were fixed in 2.5% glutaraldehyde in 0.1 M cacodylate buffer for 24 h, post-fixed in 1% osmium tetroxide, dehydrated in graded ethanol, and embedded in EPON resin. Ultrathin sections (80 nm) were stained with uranyl acetate and lead citrate and imaged using a FEI Transmission Electron Microscope at the Texas A&M Image Analysis Laboratory (RRID: SCR_022479).

### Western Blotting

Proteins were extracted using RIPA buffer containing protease/phosphatase inhibitors. Equal amounts of protein (20–40 μg) were resolved by SDS-PAGE and transferred to PVDF membranes. Membranes were probed with antibodies against METTL14 (Sigma-Aldrich HPA038002), MEETL3 (Proteintech, 15073-1-AP), WTAP (Santa Cruz sc-374280), FTO (Novus biologicals 5-2H10), vinculin (Proteintech), eGFP (Proteintech), NLRP3 (Abcam, AB270449), STING (Cell Signaling, #13647), IRF3 (Cell Signaling, #11904), pSTAT1 (Cell Signaling, #8826), STAT (Cell Signaling, #14994), LC3B (Cell Signaling, ##83506), p62 (Cell Signaling, #88588), Cleaved PARP (Cell Signaling,#5625), Gasdemin D (Cell Signaling, #97558), Caspase-8 (Cell Signaling, #4790), RIPK1 (Cell Signaling, #3493), HMGB1 (Cell Signaling, #6893). Bands were detected using chemiluminescence and quantified using ImageJ.

### qRT-PCR and RNA-seq

Total RNA was extracted using TRIzol reagent (Thermo Fisher Scientific). Complementary DNA was synthesized with the High-Capacity cDNA Reverse Transcription Kit (Applied Biosystems). Quantitative real-time PCR (qRT-PCR) was performed using SYBR Green Master Mix on a QuantStudio 7 system. Primers were designed to detect Ifnb1, Cxcl10, Ifit1, Ifit3, Oas1a, Oas1, Zbp1, with Gapdh used as endogenous controls.

For RNA sequencing, libraries were generated using the Illumina TruSeq Stranded mRNA Library Preparation Kit and sequenced on an Illumina NovaSeq 6000 platform. Sequencing reads were aligned to the mouse reference genome (mm10) using STAR, and differential expression analysis was conducted with DESeq2. Genes with a false discovery rate (FDR) < 0.05 and an absolute log₂ fold change > 1 were considered differentially expressed.

### m⁶A RNA immunoprecipitation sequencing (MeRIP-seq)

MeRIP-seq was performed using the Magna MeRIP m⁶A kit (Millipore). mRNA was fragmented to ∼100 nt and incubated with anti-m⁶A antibody. Both input and m⁶A-IP RNA were used for library preparation and sequenced on NovaSeq. Differential m⁶A peak calling was performed using exomePeak2.

## Statistics

Data are expressed as mean ± SEM. Comparisons among groups were performed using one-way ANOVA followed by Tukey’s post-hoc test. Kaplan–Meier survival curves were analyzed using the log-rank test. P < 0.05 was considered statistically significant. All analyses were performed using GraphPad Prism 10.

## Acknowledgments

We thank all the members of Xu Peng, Xiuren Zhang and Carl Tong laboratories for their helps. The work was supported by grants from R01HL145534 to C.W.T. and X. P. This material is also based upon work supported by the Texas A&M AgriLife Institute for Advancing Health Through Agriculture (IHA) and the U.S. Department of Agriculture, Agricultural Research Service, under Agreement No. 58-3091-1-018. Any opinions, findings, conclusion, or recommendations expressed in this publication are those of the author(s) and do not necessarily reflect the view of the U.S. Department of Agriculture.

